# Tadpoles of hybridising fire-bellied toads (*B. bombina* and *B. variegata*) differ in their susceptibility to predation

**DOI:** 10.1101/2020.04.02.021618

**Authors:** Radovan Smolinský, Vojtech Baláž, Beate Nürnberger

**Affiliations:** Research Facility Studenec, Institute of Vertebrate Biology, Czech Academy of Sciences, Brno, Czech Republic; Faculty of Veterinary Hygiene and Ecology, University of Veterinary and Pharmaceutical Sciences, Brno, Czech Republic

## Abstract

The role of adaptive divergence in the formation of new species has been the subject of much recent debate. The most direct evidence comes from traits that can be shown to have diverged under natural selection and that now contribute to reproductive isolation. Here, we make the case for differential adaptation of two fire-bellied toads (Anura, Bombinatoridae) to two types of aquatic habitat. *Bombina bombina* and *B. variegata* are two anciently diverged taxa that now reproduce in predator-rich ponds and ephemeral aquatic sites, respectively. Nevertheless, they hybridise extensively wherever their distribution ranges adjoin. We show in laboratory experiments that, as expected, *B. variegata* tadpoles are at relatively greater risk of predation from dragonfly larvae, even when they display a predator-induced phenotype. These tadpoles spent relatively more time swimming and so prompted more attacks from the visually hunting predators. We predict that genomic regions linked to high activity in *B. variegata* are barred from introgression into the *B. bombina* gene pool and thus contribute to gene flow barriers that keep the two taxa from merging into one.

## Introduction

Adaptation to local environments is commonplace in nature [1] and drives the evolution of novel ecotypes. Whether adaptive phenotypic divergence plays an important role in the origin of new species has been the subject of much discussion [2–7]. In this case, adaptation to different habitats should entail a marked fitness reduction for ecotypes in the ‘wrong place’ and for the hybrids between them. Of particular interest are case studies in which the selective agents and the traits under selection have been identified [7], because then the mechanisms that link adaptation to the emergence of reproductive barriers can be studied. Long-term research programmes for example on sticklebacks, sunflowers, *Heliconius* butterflies, lake whitefish and limpets have led the way in this multi-disciplinary endeavour. Relative fitnesses of different phenotypes can be estimated in different natural [8,9] or semi-natural habitats [10,11] or in experimental settings [12–14]. At the genomic level, one can infer the strength of resulting gene flow barriers from the distribution of trait effect sizes [15,16], investigate genetic linkage between traits under natural and/or sexual selection [17,18] and ask whether the inferred barrier effects are supported by patterns of introgression. Ecological divergence has long been assumed to generate barriers to gene flow in the hybrid zone of European fire-bellied toads [19], for which genomic resources have recently been developed [20]. Here we seek experimental evidence that differential adaptation of tadpoles to predators contributes to this barrier.

In contrast to the very young taxa referenced above, the fire-bellied toad (*Bombina bombina)* and the yellow-bellied toad (*B. variegata)* are more anciently derived [MRCA ∼ 3.2 mya, 20]. Nevertheless, they interbreed wherever they meet and produce a wide range of recombinants in typically narrow (2-7 km wide) hybrid zones [21]. Classic cline theory [22] suggests that the mean fitness of intermediate hybrid populations is reduced by 42% compared to that of pure populations on either side [23]. This figure presumably reflects the accumulation of diverse partial gene flow barriers since the lineage split. There is experimental evidence for intrinsic selection against hybrid embryos and tadpoles [24]. Profound ecological divergence is also strongly implicated. *B. bombina* reproduces in semi-permanent lowland ponds, such as flooded meadows and oxbows that fall dry in winter [e.g. 25]. *B. variegata* lays eggs in more ephemeral habitat (‘puddles’) and sometimes also in small steams [26], typically at higher elevations. Consequently, hybrid zones tend to be located at ecotones. Plausible arguments can be made that a large number of documented taxon differences result from adaptation to these different habitats. This applies to features ranging from anatomy over life history to the mating system [19]. However, these hypotheses need to be tested and the effects quantified to demonstrate that ecological divergence hinders gene flow.

Here we focus on tadpoles, because they experience the strongest habitat contrast. Faced with the ever present risk of desiccation, *B. variegata* tadpoles hatch from relatively larger eggs, grow faster and metamorphose earlier [27,28]. They also spend relatively more time swimming and feeding [29,30]. The less active behaviour of *B. bombina* tadpoles should protect them against visually hunting predators in ponds [29]. These taxon differences reflect a general pattern across anurans. More permanent aquatic habitats have a greater abundance and diversity of predators [31–33]. Anuran species that reproduce in them have typically quiescent tadpoles [32,34–36] that take longer to reach metamorphosis than those that breed in ephemeral sites [34,35,37]. More active tadpoles species are typically more susceptible to predation [34,38–42]. This robust pattern has been interpreted as a trade-off [34,37,43,44]: tadpoles can either be highly active and maximise food intake for fast growth and development in temporary habitats or avoid predators through quiescent behaviour in more permanent sites, but not both. Tadpole activity and growth rate covary as expected in evolutionary contrasts [45] but evidence for a causal relationship is mixed at the intra-specific level [46–51]. However, the trade-off between rapid growth and predator avoidance is almost universal [52] irrespective of whether activity is the mechanism that links the two. We therefore expect that *B. variegata* tadpoles are at greater risk of predation than *B. bombina*. Any traits associated with such increased risk would therefore be unlikely to introgress into the *B. bombina* gene pool and so represent a partial barrier to gene flow between the taxa.

Nevertheless, *B. variegata* tadpoles do sporadically encounter predators such as newts, dragonfly and dytiscid larvae, water scorpions and salamander larvae [26]. Thus, tadpoles would maximise their chance of reaching metamorphosis by adjusting their phenotype to the local balance of risk. Indeed, a wide range of anurans display phenotypic plasticity in response to predators. Almost invariably, tadpoles reduce their activity level when they perceive chemical cues that indicate predator presence [37,53,54]. This response can occur within minutes [55]. Longer term exposure to such cues can cause an increase in refuge use [56,57], alter larval morphology [54], prompt the development of colour spots on the tail [58,59], and slow growth and development [60,61]. These effects are seen across venues from the laboratory to the field [62,63].

The commonest and most closely analysed morphological response to predators is the development of a relatively deeper tail fin [35,36,47,54], usually in combination with a shorter tail and a shallower body. A deeper tail may facilitate rapid escape (faster burst swimming speed) or lure attacks away from the more vulnerable body. While there is evidence that natural variation in tail shape affects tadpole swimming performance [42,64], studies to date have given stronger support to the lure hypothesis [65,66]. Irrespective of the particular mechanism, there is evidence that predator-induced phenotypes are eaten less than predator-naive ones [67–70] and that survival is correlated with measured traits that show a plastic response [70–72]. A recent study of *Rana temporaria* [73] makes a particularly strong case for adaptive phenotypic plasticity in both behaviour and morphology, including an assessment of the variability in predation risk within and between ponds and over time that could bring about such a flexible strategy.

Both *B. bombina* and *B. variegata* respond to the presence of predators with reduced activity [29,30] and by developing a relatively deeper tail fin [74, see also 54]. Exposure to chemical predation cues slows their growth and development under laboratory conditions [74]. At present, we do not know whether the induced phenotypes offer better protection from predators. Because we want to know whether *B. variegata* would suffer higher predation than *B. bombina* in ponds, we contrast induced phenotypes only. Two previous studies addressed the relative predation risk of these toads. In a comparison of predator-naïve tadpoles, *B. variegata* suffered relatively higher predation from a range of predators *[Aeshna, Dytiscus, Triturus;* 29]. In the second study with predator-induced tadpoles, a strong population effect overwhelmed any taxon differences [two populations per taxon; 74] In both cases, significant effects were observed in the first half of the larval period. Observations in outdoor mesocosms show that the overall rate of predation by dragonfly larvae is highest during that time and drops off sharply afterwards [75]. Similarly declining rates of dragonfly predation have been reported for two species of *Ran*a [76].

Here, we estimate the relative susceptibility to predation of pure *B. bombina* and *B. variegata* throughout the first four weeks post-hatching, i.e. until the first *B. variegata* tadpoles reach metamorphosis. Dragonfly larvae were used as predators, because they are ubiquitous in ponds and also found in puddles. We also assess the contributions of activity, body size and tail shape to the observed differences in predation risk.

## Materials and Methods

### Animal collections, pairing scheme and toad care

Adult toads were collected from Moravia (Czech Republic) where the Carpathian lineage of *B. variegata* meets the Southern lineage of *B. bombina* [77]. The experiment was carried out in 2017. Starting in early May, aquatic habitats were visited regularly (14 known *B. bombina* sites and 19 known *B. variegata* sites). Due to an unusually cold spring, toads did not appear in appreciable numbers until early June and were then promptly collected to ensure that females had not yet laid eggs. Collection sites (Table 1) were located at least 6 km from known areas of hybridisation and deemed ‘pure’ based on the taxon-specific spot pattern on the toads’ bellies [19]. For each taxon, minimum and maximum distances between sites were 1.5 and 43.6 km (*B. bombina*) and 0.7 and 14.9 km (*B. variegata*). *B. bombina* sites were ponds with a minimum width of 5 m and a maximum depth greater than 1 m. They all contained predators (incl. *Aeshna cyanea, Natrix natrix*, fish) and, with one exception (site E), they had abundant aquatic vegetation. *B. variegata* sites were either water-filled wheel ruts (max. depth < 30 cm) or in one case (site 2) the shallow end of an abandoned swimming pool. They were devoid of aquatic vegetation and three of nine sites contained no predators at all (see S1 Table for details).

**Table 1.**
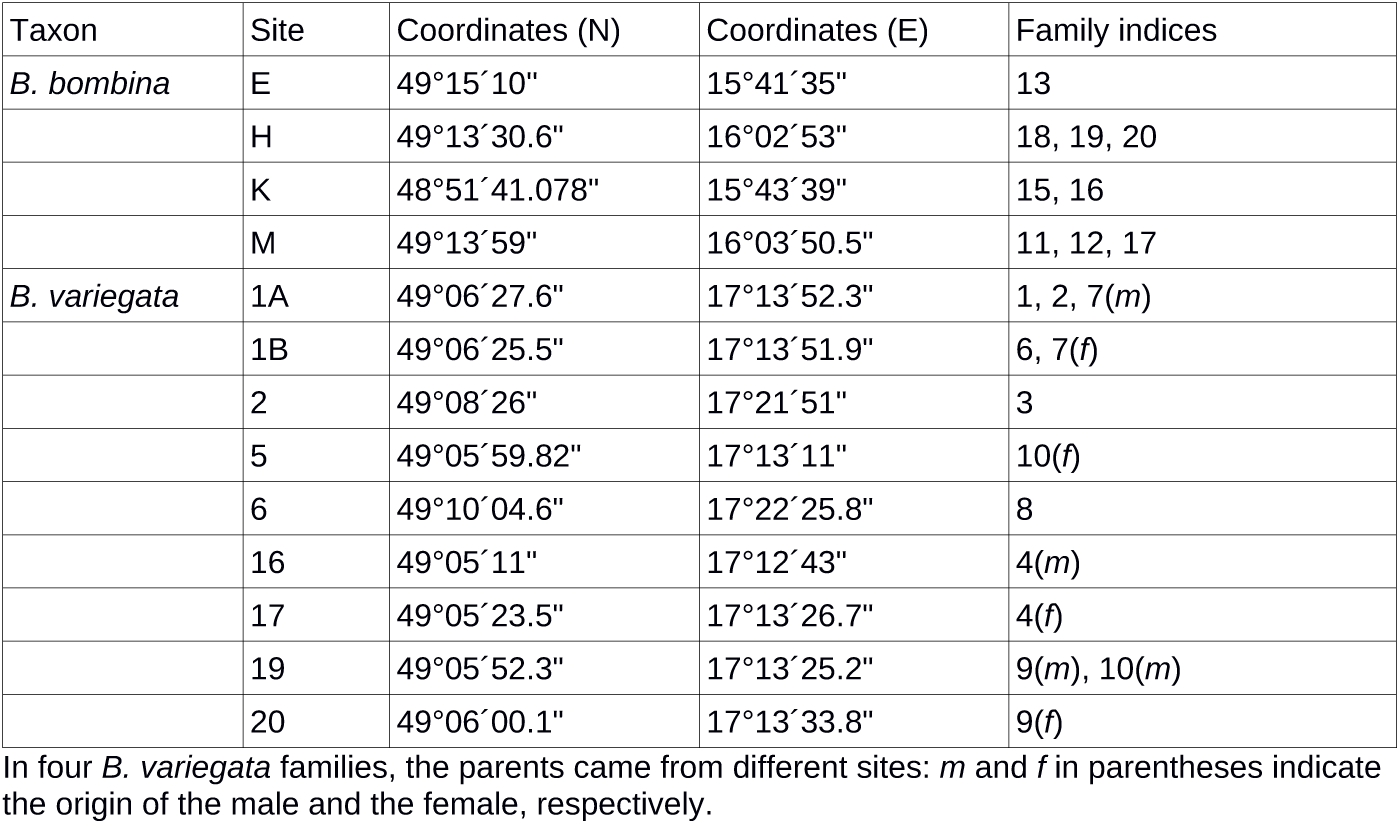
Collection sites and pairing scheme.

| Taxon | Site | Coordinates (N) | Coordinates (E) | Family indices |
| --- | --- | --- | --- | --- |
| *B. bombina* | E | 49°15'10" | 15°41'35" | 13 |
|  | H | 49°13'30.6" | 16°02'53" | 18, 19, 20 |
|  | K | 48°51'41.078" | 15°43'39" | 15, 16 |
|  | M | 49°13'59" | 16°03'50.5" | 11, 12, 17 |
| *B. variegata* | 1A | 49°06'27.6" | 17°13'52.3" | 1, 2, 7(*m*) |
|  | 1B | 49°06'25.5" | 17°13'51.9" | 6, 7(*f*) |
|  | 2 | 49°08'26" | 17°21'51" | 3 |
|  | 5 | 49°05'59.82" | 17°13'11" | 10(*f*) |
|  | 6 | 49°10'04.6" | 17°22'25.8" | 8 |
|  | 16 | 49°05'11" | 17°12'43" | 4(*m*) |
|  | 17 | 49°05'23.5" | 17°13'26.7" | 4(*f*) |
|  | 19 | 49°05'52.3" | 17°13'25.2" | 9(*m*), 10(*m*) |
|  | 20 | 49°06'00.1" | 17°13'33.8" | 9(*f*) |

In four *B. variegata* families, the parents came from different sites: *m* and *f* in parentheses indicate the origin of the male and the female, respectively.

Males and females were paired within sites whenever possible. However due to unequal sex ratios per site, four *B. variegata* pairs involved toads from separate, but nearby sites (Table 1, maximum distance between sites per pair: 0.96 km). Each pair was housed in a 52 x 35 cm plastic box with 7L of well water, a polystyrene island and three pieces of plastic aquarium plants as support for egg batches. As a precaution against the potential spread of *Batrachochytrium* infection, all waste water from the entire experiment was heated to 100° C and treated with UV (Eheim UV reflex UV800) before disposal to kill any fungal spores.

On 20 June, a belly swab was taken from each toad to test for *Batrachochytrium dendrobatidis (Bden)* and *B. salamandrivorens (Bsal)* infection. The former pathogen is widely distributed in the Czech Republic, but so far no population declines have been observed [78]. The qPCR followed the protocol of Blooi et al. [79] using a Roche Light Cycler480. No evidence of either infection was found. We note that samples in the same essay but from an unrelated study scored positive for *Bden* infection. The adults were subsequently returned to their exact collection sites.

### Predators

We used *Aeshna cyanea* larvae as predators. This dragonfly is widely distributed across Europe and was present in most of the habitats we surveyed (see above). Two hundred larvae were collected from one site near Jihlava (Vysočina County) and were kept in groups of 15 in 50L plastic boxes. They were fed every other day with Tubifex worms before the predation trials started and with tadpoles thereafter.

### Oviposition

To induce reproduction, 250 I.U. of human chorionic gonadotropin, dissolved in 0.2 ml of amphibian Ringer solution, were injected into the lymph sac of an outer thigh of both males and females on 6 June. Most pairs produced egg clutches overnight. Over the following two days, 30-40 eggs per family were transferred into the rearing set-up (see below). The remaining offspring were raised in the oviposition boxes, with weekly water changes. They were used as food for the dragonflies in order to condition the water in the rearing set-up. The adults were moved to new boxes of the same size. Tadpoles hatched on 12-13 June. In the following, we refer to 13 June as day 0 of the experiment.

### Tadpole rearing

We constructed two gravity flow circuits. For each of them, five plastic tubs (52 x 37 x 16 cm) were interconnected with silicone tubing such that water could be pumped from a 20 L bucket at floor level into the top tub and then descend stepwise via the other four tubs before returning to the bucket. Inflows and outflows in each tub were positioned diagonally opposite to maximise mixing. Each tub held 27L of water at a depth of 14 cm. In order to maintain uniform water quality, the pumps (Eheim StreamON+ 2000) were operated 24 hours per day at a low flow rate and per circuit 7 L of water were replaced daily. Throughout the experiment, water temperature was 22-24°C and the air temperature was 24-26°C. All water changes for adults and tadpoles were carried out with well water heated to 22°C and sterilised with UV (Eheim UV reflex UV800).

The upper four tubs in each circuit were subdivided with fibreglass insect mesh into eight compartments (size: 18 x 11.5 x 16.5 cm, 64 compartments in total). A plastic aquarium plant was added to each compartment for structural complexity. At the start of the experiment, each compartment held 10 hatchlings from a single family. Families (nine per taxon) were distributed over either three or four compartments in total and either one or two compartments in each circuit. Exactly half of the compartments in each circuit were assigned to each taxon. Placement of families within circuits was decided by random draws.

Tadpoles were fed boiled nettle (*Urtica dioica*) leaves *ad libitum* and uneaten food was removed every day. In order to simulate predator presence, we let chemical cues of tadpole predation circulate from day 0 onwards. For this, we used the fifth tub in each circuit to feed 1-2 non-experimental tadpoles (depending on size) to one dragonfly larva every other day.

### Predation trials

Predation trials started on day 2 when the tadpoles were active. They were carried out in two plastic boxes subdivided with fibreglass insect mesh into four compartments (26 x 26 x 18 cm), each with two plastic aquarium plants (akin to *Myriophyllum*). They contained a 1:1 mixture of well water and water from the rearing circuits. For each trial, four tadpoles (2 *B. bombina*, 2 *B. variegata*) were placed into a given compartment and allowed to acclimate for 30 min. A dragonfly that had not been fed on the previous day was then added and the trial was stopped when the first tadpole had been eaten (maximum duration: 45 min). The identity of the missing tadpole was determined from digital photographs of the animals taken before the trial (see below). The survivors were returned to their rearing compartments. The trials were recorded with a Microsoft LifeCam VX-2000 video camera suspended over each experiment tub.

Predation trials were carried out on 18 days as follows: two consecutive experiment days were followed by one ‘off-day’ (1 exception); this pattern was repeated nine times until day 29 post hatching, when the first *B. variegata* tadpoles entered metamorphosis. There were 20 trials per day between 10:00 and 15:00 hours. A custom Perl script was used to haphazardly draw the composition of each trial per day such that (a) four different families were represented, (b) one tadpole per taxon came from circuit 1 and the other from circuit 2 and (c) each rearing compartment contributed at least one tadpole on a given day. However towards the end of the experiment, a few rearing compartments were empty and the contribution of others had to be increased accordingly. No individual tadpole or dragonfly was used more than once per experiment day.

### Morphometrics

Digital images were taken by placing a tadpole into a glass cuvette and photographing it from the side with a Canon EOS 400D camera (lens: Canon EF 28-90mm). TPS Dig (Software by F. James Rohlf) software was used to take the following measurements from these images: total length, tail length, body height and maximum tail height.

### Behaviour

From the video footage of the predation trials, we recorded the behaviour of a focal tadpole during one 30 sec interval per trial for half of the trials on a given day. A custom Perl script provided the pseudo-random start point of the observation interval and allowed behaviour (swimming, feeding, resting) to be recorded with separate key strokes. From one trial to the next, we alternated between a *B. bombina* and *B. variegata* tadpole. Taxon identification was based on size in the early trials. Morphological differences were used as soon as they became apparent. Footage from the first two days was not used, because the taxa could not be reliably distinguished. We discarded all observation intervals that overlapped with a predator attack. We also recorded for 68 predator attacks (four on each on 17 experiment days) whether they were initiated by and directed towards an actively swimming tadpole.

Collection permits were granted by the relevant regional offices (South Moravia and Zlín) of the Czech Ministry for the Environment. The experimental protocol was approved by the ethics committee of the Institute of Vertebrate Biology, CAS, permit no. 15/2016.

## Statistical analyses

### Estimates of predation risk

We estimated predation risk per rearing compartment as the number of tadpoles eaten divided by the number of valid trials in which that compartment was represented. Especially with a small number of trials, this estimate may be biased if tadpoles from a certain compartment happen to be assigned to trials with, say, particularly well defended congeners. There is a range of statistical methods to estimate an individual’s skill from a number of contests with opponents of varying strength (as e.g. in chess), starting with the model of Bradley and Terry [80]. However, none of these match our trial design of a single winner (or rather: looser) and a ‘tie’ for the other three. For each rearing compartment, we therefore computed the expected ‘risk context’ from a large number of simulated trials and compared this estimate with the mean risk context in the actual trials. A close match indicates the absence of bias. Risk context is here defined as the mean predation risk of the other three rearing compartments represented per trial, averaged over all trials in which the focal compartment was represented. Using the same custom script as before, we simulated 2550 trials and used these to compute the expected risk context of all 64 rearing compartments.

We investigated the effects of taxon, circuit and family on predation risk per rearing compartment. Circuit was treated as a fixed effect, because it has only two levels. Family was modeled as a random effect nested within taxon. We did not include a population term in the model, because populations were unevenly represented and some families had parents from different sites. Any differences between populations should appear as family effects. Linear mixed models were fitted with the R module *lme4* [81]. Model comparisons were carried out with likelihood ratio tests.

### Morphological correlates of survival

Logistic models were used to investigate the effects of taxon, tadpole size and tail shape on survival per trial. The input was generated by sorting all valid trials in chronological order and picking from them alternately (a) a surviving tadpole at random or (b) the non-surviving tadpole. For the entire dataset, the four morphological measurements were log-transformed to improve normality and entered into a principal component analysis (PCA). We treat the first principal component as a measure of tadpole size. From a regression of log(tail height) on PC1 we extracted the residuals as a measure of relative tail height (≈ tail shape). To ensure shared allometry between the taxa, we inspected pairwise plots of the four measured variables [82]. For the focal tadpole per trial, PC1 and residual log(tail height) were expressed as a deviation from the trial mean. All analyses were carried out in R v.3.4.4.

## Results

### Taxon differences in predation risk

In total, there were 360 trials. Of these, no tadpole was eaten in 96 trials, more than one was eaten in five trials and in four trials the two taxa were not evenly represented. Discarding these, 256 valid predation trials remain. The proportion of valid trials in which a *B. variegata* tadpole was eaten was 0.65. There was also a slightly higher risk for tadpoles from circuit 1 as opposed to circuit 2 (proportion = 0.56). This effect was largely restricted to *B. bombina* (see below). The number of trials in which no tadpole was eaten was larger in the second half of the experiment (tadpole age: 16-29 days, n = 66) than in the first half (tadpole age: 2 - 15 days, n = 30).

Predation risk was computed per rearing compartment as the number of tadpoles eaten during valid predation trials over the number valid trails in which this compartment was represented. The observed risk context (as defined in the Materials and Methods) closely matched the prediction from simulated trials (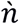per compartment = 159.9): for 56 of 64 compartments the observation deviated by less than 10% from the expectation (max. deviation 14%) and in all cases it was well within one standard deviation of the prediction (S1 Fig). We therefore use predation risk per rearing compartment as the response variable in the following analyses. The estimates are plotted in Fig 1.

**Fig 1.**
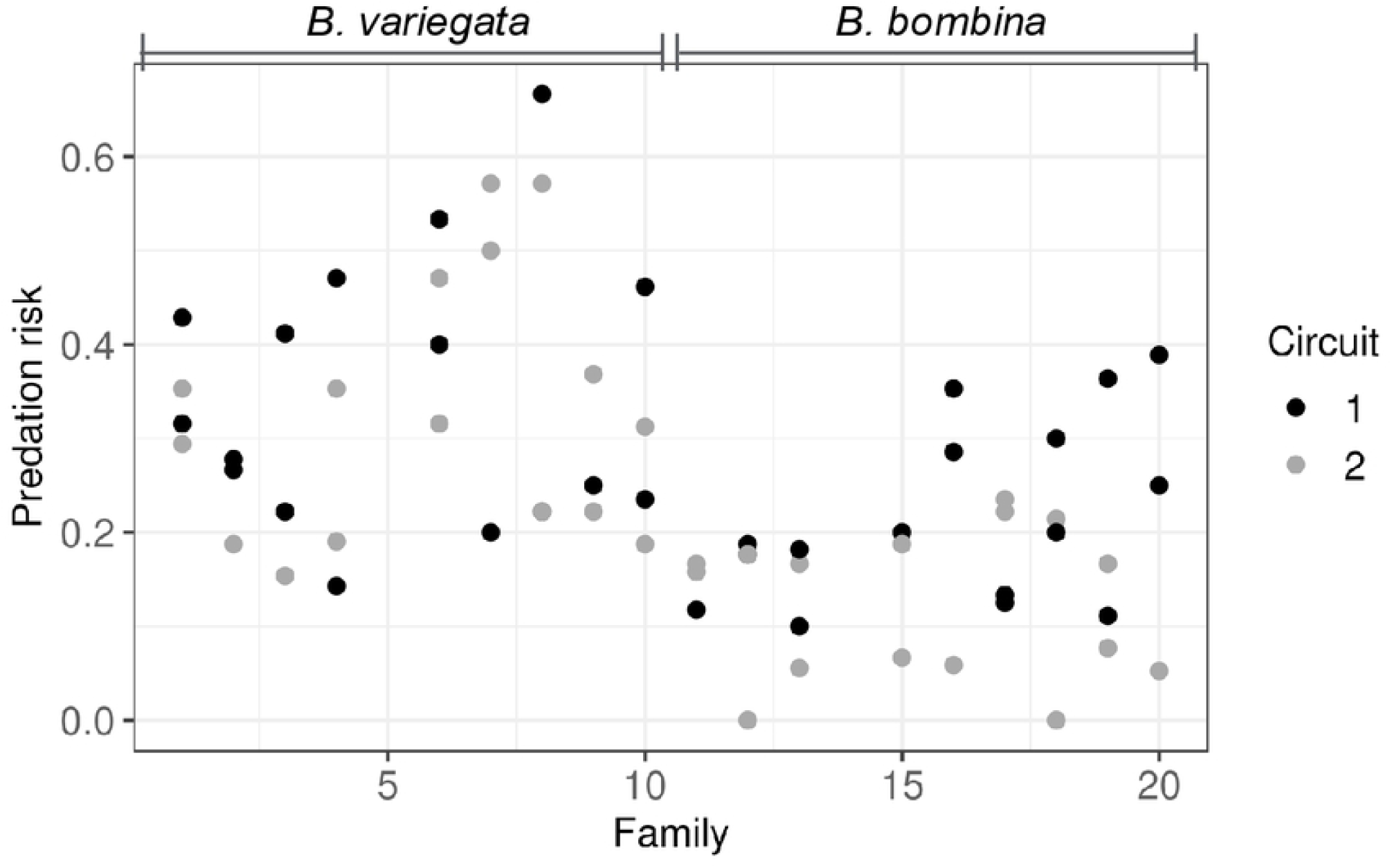
Predation risk per rearing compartment arrayed by family. *B. variegata*: families 1-4 and 6-10, *B. bombina*: families 11-13 and 15 - 20.

The full linear mixed model (Table 2, model 1) included taxon, family (nested within taxon) and circuit as main effects and also interactions between taxon and circuit and between family and circuit. Model comparison showed that neither of interaction terms were significant (models 2 *vs* 1: p = 0.0829; models 3 *vs* 1: p = 1) and so both were dropped (model 4). The main effect of taxon was overwhelmingly significant (models 5 *vs* 4: p = 1.911e-05), providing clear evidence that *B. variegata* tadpoles were at relatively greater risk of predation by *Aeshna cyanea* larvae. There was also a significant circuit effect (models 6 *vs* 4: p = 0.0278). Zero variance was assigned to the family effect in models 3 - 6. Two-way ANOVAs of taxon and circuit closely reproduced the patterns of significance for these two factors (analyses not shown). While there is clear evidence for a taxon effect and a minor effect of circuit, a large amount of unexplained variation remains (Fig 1).

**Table 2.**
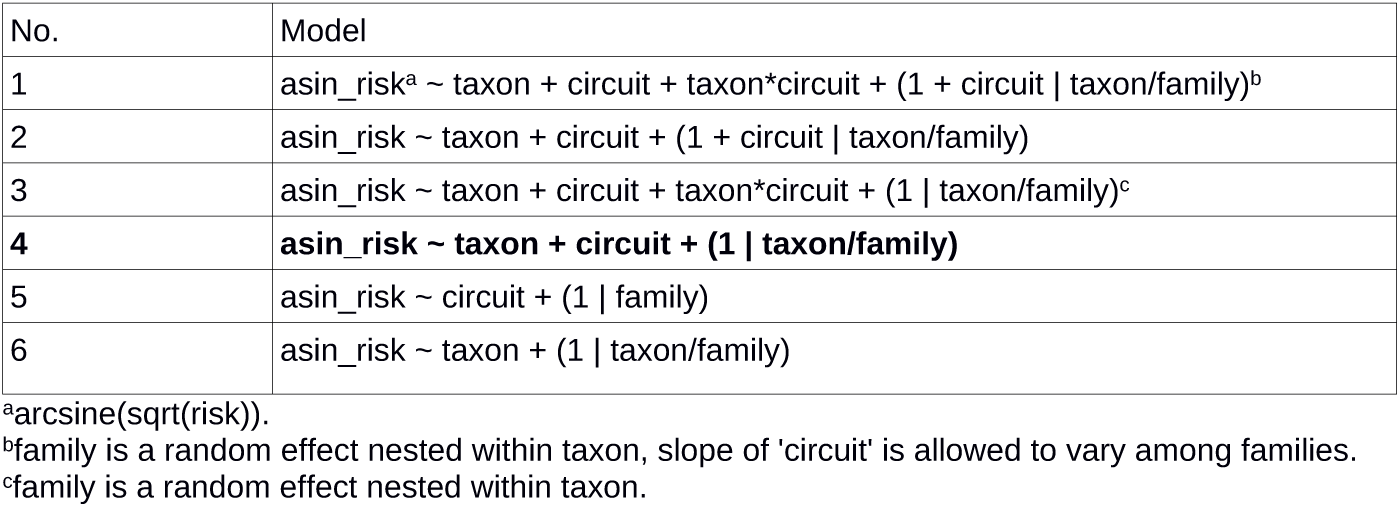
Linear mixed models of predation risk per rearing compartment.

### Tadpole size and shape

Throughout the experiment, *B. bombina* was on average smaller with a shallower tail (Figs 2A and 1B). But after correction for body size, *B. bombina’s* tail fin was relatively higher than that of *B. variegata* (Fig 2C). We note that our shearing approach to correct for body size is validated by the shared allometry in both taxa (S2 Fig). If size and/or tail shape are functionally important, we would expect to see within-taxon correlations with survival as well. The logistic regression analysis (Table 3, see Materials and Methods for details) confirmed the significantly lower survival of *B. variegata* and also showed a strong interaction of tadpole size (PC1) and taxon. There were no significant effects associated with relative tail height.

**Table 3.**
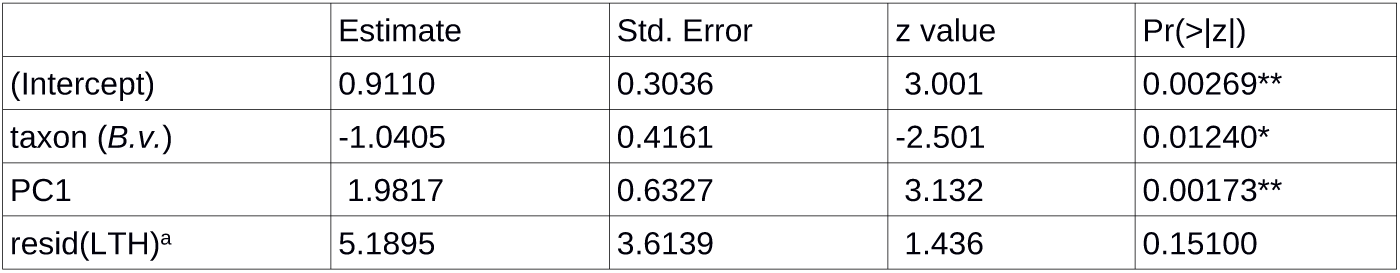
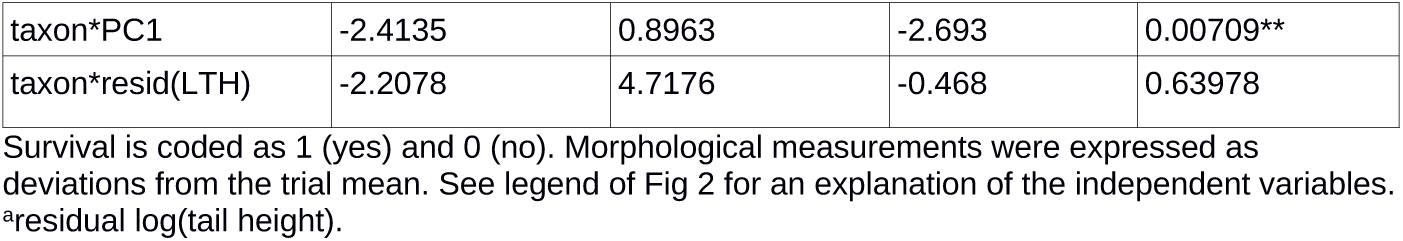
Coefficients of the logistic model of survival.

**Fig 2.**
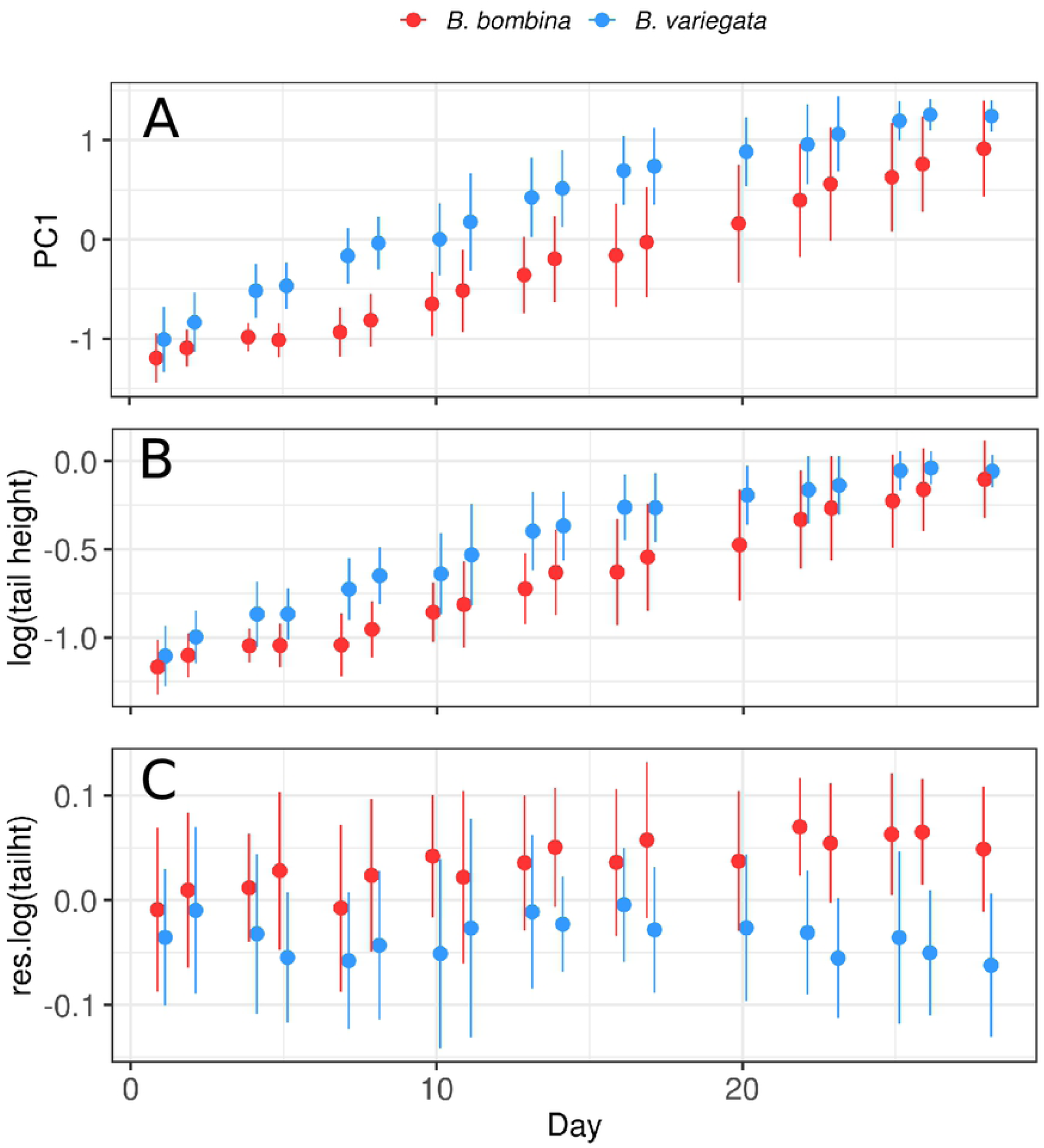
Morphological development by taxon. Error bars represent one standard deviation. PC1 = 1. principal component ≈ body size, res.log(tailht) = residual log(tail height) ≈ size-corrected tail height.

Fig 3A illustrates that larger *B. bombina* tadpoles survived better than smaller ones (t = 3.5993, df = 85.589, p-value = 0.0005). But this effect is not consistent across the whole dataset. *B. variegata* tadpoles are overall larger and more at risk of predation, whereas smaller size is associated with greater risk in *B. bombina*. There is therefore no evidence that a general preference of *Aeshna* larvae for larger prey causes higher mortality in *B. variegata* tadpoles. The analogous plot for relative tail height hints at an advantage for higher tail fins (Fig 3B), but the present dataset provides no statistical support for this hypothesis.

**Fig 3.**
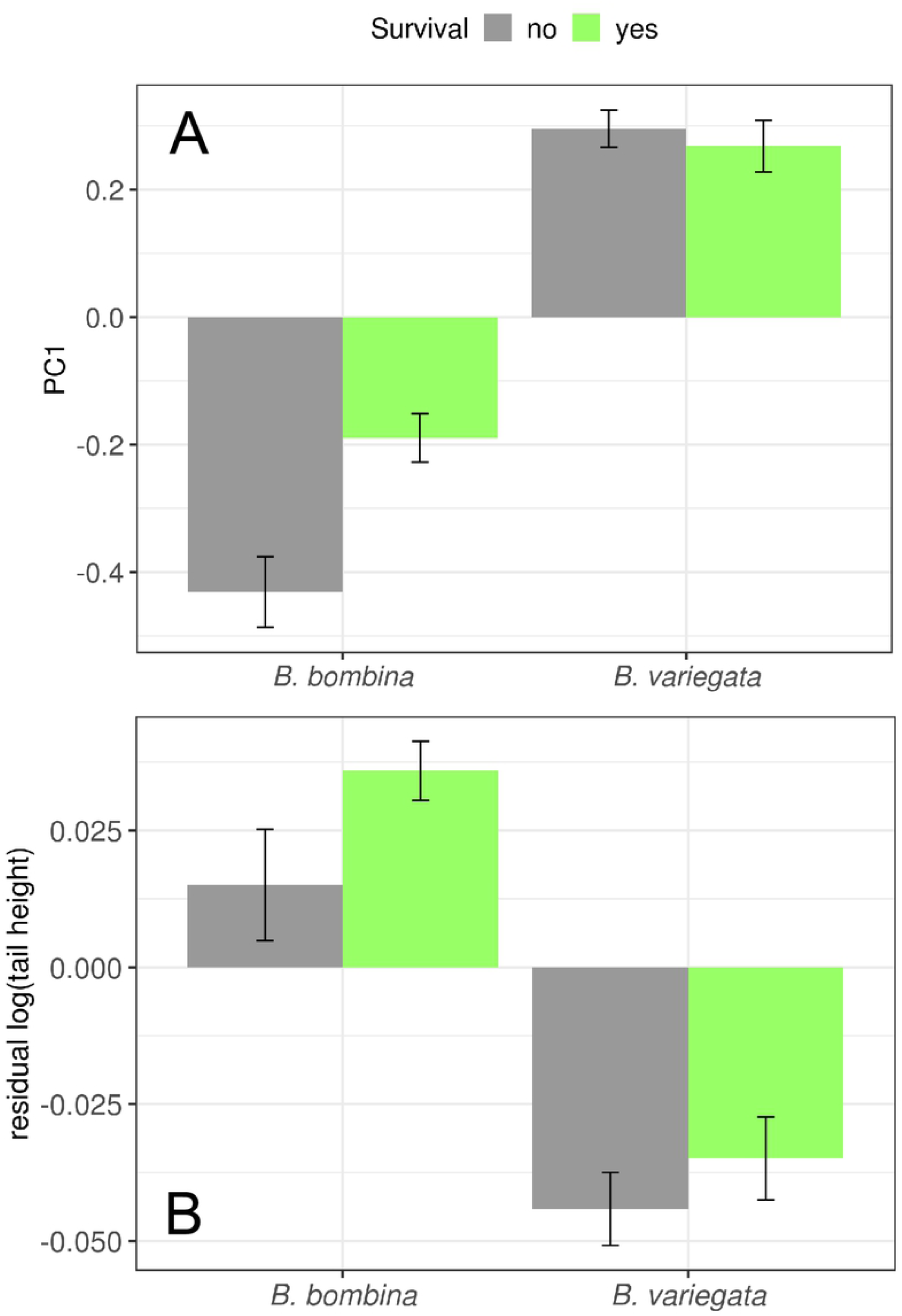
Means of body size and tail shape by taxon and survival. Plotted are the means from the logistic regression dataset for PC1 (A) and residual log(tail height) (B). Error bars indicate one standard error of the mean.

### Tadpole activity

During the 30 sec observation intervals, both taxa spent most of their time either resting or, seemingly, feeding. No food was provided in these trials. So, the tadpoles’ persistent scraping of surfaces with their mouthparts, especially during the first half of the experiment, probably provided little, if any nutrition. Swimming typically occurred in short bursts of a few seconds separated by inactive intervals. Ninety-two percent of recorded predator attacks were directed at actively swimming tadpoles.

The distributions of the time spent swimming were strongly skewed to the right in both taxa (Fig 4). As expected, *B. variegata* spends more time swimming than *B. bombina* (means: 4.60 and 1.75 sec, respectively; unpaired Wilcoxon rank sum test, W = 2234.5, p-value = 0.0096). Focal tadpoles were chosen based on their clear visibility during the 30 sec observation interval. Their survival status is unknown. We can therefore not investigate within-taxon correlations between activity and survival.

**Fig 4.**
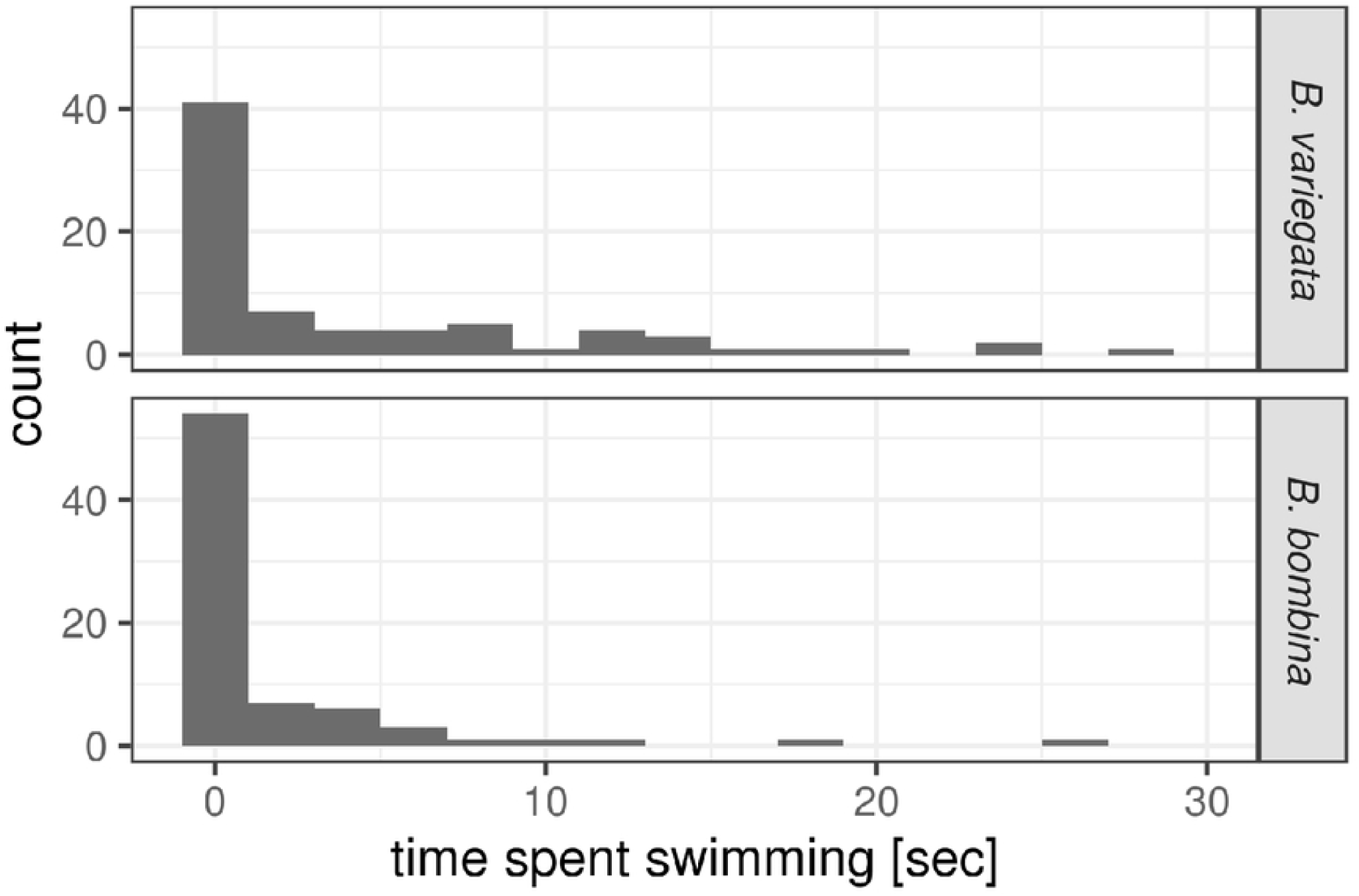
Time spent swimming during 30 sec observation intervals by taxon. Number of intervals: 76 and 75 for *B. variegata* and *B. bombina*, respectively.

### Predation rates over time

A logistic model with taxon and experiment day as independent variables produced only a non-significant taxon x day interaction (analyses not shown). There is thus no indication that the relative survival of the two taxa changed over the course of the experiment. Fig 5 illustrates the proportion of *B. variegata* tadpoles consumed and the number of valid trials over time. Not only were there fewer valid trials in the second half of the experiment (tadpole age: 16-29 days), the mean duration of trials also increased from 9.7 min to 13.6 min.

**Fig 5.**
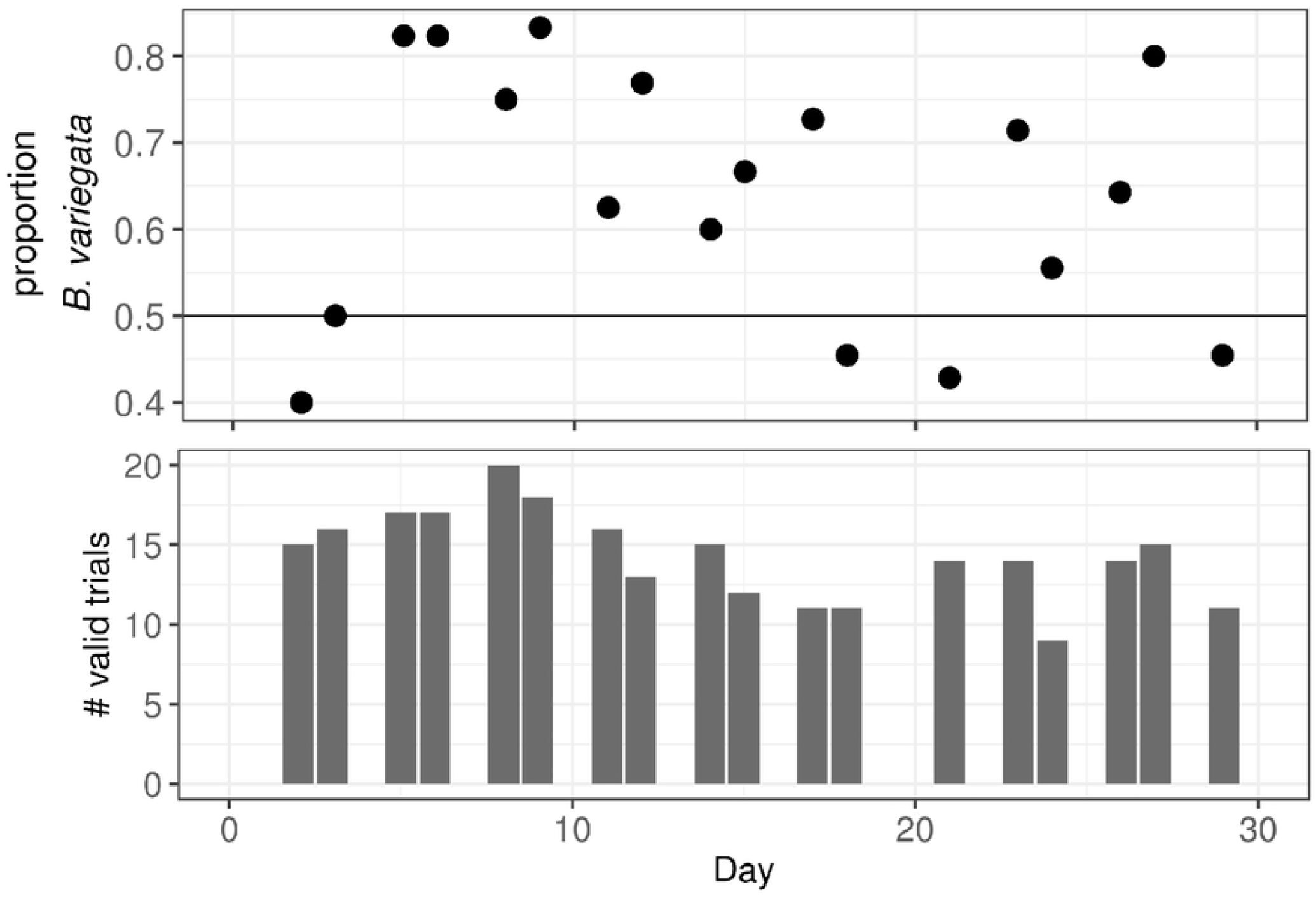
Relative predation risk and number of valid trials per day. A. The proportion of tadpoles consumed that were *B. variegata*. B. The number of valid trials per experiment day (out of 20).

## Discussion

In our experimental setting, tadpoles of the puddle breeder *B. variegata* are at greater risk of predation from dragonfly larvae than those of *B. bombina* that develop in semi-permanent ponds. This result confirms our expectation based on similar species comparisons [35]. It also agrees with the findings of Kruuk and Gilchrist [29] who conducted predation trials with larger groups of predator-naive *B. bombina* and *B. variegata* tadpoles that could only adjust their behaviour. Here, we allowed tadpoles to develop a more complete predator-induced phenotype. Nonetheless, the survival difference between the taxa persisted and it was particularly strong during the first two weeks of larval development (Fig 5), when risk from dragonfly predation is highest [75]. Vorndran et al. [74] also reared tadpoles in the presence of chemical predation cues and found a marked survival difference between tadpoles from the two replicate *B. variegata* populations. The two collection sites in Romania also differed in predator abundance and proximity to the hybrid zone, observations that warrant more detailed population comparisons in that area [83].

Before we interpret our findings more fully, we consider potential limitations of our experimental design. Many tadpoles were used in more than one trial. Later trials may therefore have involved individuals that benefited from experience gained and/or were inherently better at avoiding predators. We believe that this resembles the situation in nature more closely than the alternative of shielding tadpoles from predator encounters until the day of their one experimental test. As pointed out by Melvin and Houlahan [84], typical tadpole mortality in nature (approx. 95% according to their review) may be non-random with regard to traits under study. The typically much more benign conditions in the laboratory may thus produce experimental cohorts that are not representative of the population of origin. While we have not reproduced the selective regime of any semi-permanent pond, repeated exposure to dragonfly larvae adds realism to our study. Nevertheless, we consider this close-up view of predator-prey interactions in the laboratory as a piece in a mosaic that should be complemented with experiments in semi-natural settings and using additional predator species.

If *B. bombina* and *B. variegata* were reproductively isolated, these results would address the question why *B. variegata* reproduces only rarely in semi-permanent, predator-rich ponds [85]. Whether or not *B. bombina* tadpoles are present in these habitats may not matter, because there is little evidence for competition among tadpoles in ponds [33,35,38]. Predators tend to keep tadpole numbers sufficiently low so that competition is negligible. *B. bombina* then simply provides a benchmark for a successful ‘pond strategy’. *B. variegata*’s greater susceptibility to predation shows that its tadpoles fall short of this benchmark even when they display the predator induced phenotype. Other traits such as relatively fewer eggs per clutch [27] may also limit *B. variegata’s* reproductive success in ponds.

Hybridisation shifts the focus away from a comparison between distinct types and towards selection on traits in recombinants. These genotypes predominate in *Bombina* hybrid zones, while F1 individuals are very rare, if present at all [21,23]. Each generation, associations between traits are broken up by hybrid matings but also replenished by the influx of pure types from the periphery. Theory predicts a dynamic equilibrium between these processes that determines how strongly the hybrid zone functions as a genetic barrier to gene flow [86,87]. Any trait that increases the susceptibility of tadpoles to predation is unlikely to introgress into the *B. bombina* genepool and it should impede the introgression of traits with which it is genetically linked. In this way, traits that mediate adaptation to either ponds or puddles contribute to partial reproductive isolation in *Bombina*.

We considered traits that are expected to affect the risk of being detected by predators and the ability to escape once attacked. Prey activity alerts visually hunting predators and correlates strongly with tadpole predation rates in inter-specific comparisons (see Introduction) and also within species [67,88]. In our experiments, *B. variegata* spent on average more time swimming than *B. bombina*. Since swimming was the single most important trigger of dragonfly attacks, activity contributes importantly to the difference in predation rate between the taxa. There was no evidence that relatively higher tail fins caused better survival in either taxon. The large variances that rendered differences between group means (taxon, survival) non-significant (Figs 2C and 3B) may reflect heritable variation among families in the amplitude and type of morphological plasticity [89]. This trait warrants further study using more detailed morphometrics [64]. Finally, our analyses disproved the hypothesis that an overall preference of *Aeshna cyanea* larvae towards larger prey contributed to *B. variegata’s* greater mortality.

Is higher predation risk in *B. variegata* a direct consequence of adaptation to ephemeral habitat? Population studies have shown that successful metamorphosis can be restricted in a given year to less than 50% of sites in which eggs have been laid (depending on annual rainfall) and that desiccation is the main cause of reproductive failure [90–92]. This is reflected in the growth rate of *B. variegata* tadpoles which was the fastest in a set of 15 European and NE North American anurans [60]. Intra-specific competition should be commonplace as tadpoles grow and puddles shrink [93]. The high activity of tadpoles growing in this type of habitat supports this [94]. In the lab, large *B. variegata* tadpoles displace smaller conspecifics from food patches with their powerful tail movements [95]. If indeed *B. variegata’s* larval development has been mainly shaped by time constraints and competition, there may be trade-offs that limit the effectiveness of anti-predator strategies. Indeed predator-induced plasticity in *Bombina* entails slower growth and delayed metmorphosis [74]. Based on studies with *Rana* tadpoles [96,97], we expect all of these responses to be mediated by the neuroendocrine stress axis and by levels of corticosterone in particular.

While phenotypic plasticity involves the same traits in both taxa, the differences between the induced phenotypes are not just a matter of degree. There are qualitative differences as in the body size effect on predation risk that is present in *B. bombina* but not in *B. variegata* (see above). Moreover in *B. variegata*, absolute tail height shows no response to predator presence. The increase in relative tail height reflects reduced growth rates of other body parts. In contrast, all measured dimensions showed reduced growth in *B. bombina*, with an allometric change in tail height [74]. Finally, *B. variegata* frequently develops dark spots on the tail fin that may divert attacks away from the body, whereas *B. bombina*’s tail fin is always uncoloured [98]. Assuming that tail spots provide protection, the genomic regions linked to them may even show adaptive introgression. As *B. bombina* and *B. variegata* adapt to different breeding habitats, they may evolve somewhat different strategies to counter common selection pressures [99]. This could reduce the fitness of hybrids in which these strategies are broken up. Based on this study, we expect that higher activity in *B. variegata* tadpoles acts as a barrier to gene flow, but other ‘component traits’ of predation risk may also be under selection in the hybrid zone und some may be conduits rather than barriers of gene flow.

## Acknowledgments

We thank Oliver Gast and Zuzana Hiadlovská for help in the field and with the experiments, respectively. Jarrod Hadfield, Lumír Gvoždík and Crispin Jordan offered advice on the experimental design and statistics. We thank Günther Gollmann, Jacek M. Szymura and Josh Van Buskirk for comments on an earlier version of this manuscript.

## Supporting Information

**S1 Fig. Risk context per rearing compartment.**

**S2 Fig. Pairwise plots of morphological traits.**

**S1 Table. Ecological features of the collection sites.**

